# Unconventional myosin VI is involved in regulation of muscle energy metabolism

**DOI:** 10.1101/2025.05.13.653637

**Authors:** Dominika Wojton, Dorota Dymkowska, Damian Matysniak, Malgorzata Topolewska, Maria Jolanta Redowicz, Lilya Lehka

**Affiliations:** Laboratory of Molecular Basis of Cell Motility, Nencki Institute of Experimental Biology, Polish Academy of Sciences, 3 Pasteur St., 02-093 Warsaw, Poland; Laboratory of Cellular Metabolism, Nencki Institute of Experimental Biology, Polish Academy of Sciences, 3 Pasteur St., 02-093 Warsaw, Poland

**Keywords:** Energy metabolism, mitochondria, muscle fiber types, myogenic cell, skeletal muscle, unconventional myosin VI

## Abstract

Mitochondria are essential for regulation of the metabolic state of skeletal muscle, making their structure and function crucial for muscle performance. Myosin VI (MVI), an unconventional minus-end-directed motor, is expressed in skeletal muscle and myogenic cells. To explore its role in mitochondrial function and muscle metabolism, we used MVI knockout mice (*Snell’s waltzer, SV*) and their heterozygous littermates. We analyzed muscle samples from newborn (P0) and adult mice (3- and 12-month-old) and found that both MVI mRNA and protein levels were highest in newborn muscles and decreased with age. MVI expression also varied by muscle type, being highest in the slow-twitch soleus muscle (SOL) of adult mice. Loss of MVI had the most significant effects on SOL, which contains the highest number of mitochondria compared to fast-twitch muscles. MVI loss resulted in reduced respiratory capacity and ATP production in myogenic cells, indicating impaired mitochondrial function. Furthermore, MVI deficiency caused a shift from glycolytic to oxidative fiber types, especially in SOL. We also observed increased phospho-AMPK levels in MVI-KO SOL across all time points, along with downregulation of the mTOR pathway and upregulation of proteins involved in lipolysis. These findings highlight MVI as a novel regulator of metabolic processes in skeletal muscle.

## 1. Introduction

Myosins form a large superfamily of motor proteins relying on the energy from ATP hydrolysis to move along the actin tracks. Engaged in a multitude of cellular functions, myosins are categorized into various classes. Among them, the muscle myosin isoforms are the most renowned and extensively studied, being pivotal in orchestrating muscle contraction. Class II myosins, encompassing both muscle and non-muscle myosins (structurally and functionally resembling muscle isoforms, NMII) form the most prevalent subgroup within the myosin superfamily [^1^]. Myosins II, encoded by *MYH* genes, are also termed conventional ones as they form large bipolar filaments via tail-directed oligomerization. All other myosins are termed as unconventional (UMs) since they are not able to form filaments, although some can dimerize; UMs are encoded by *MYO* genes. Several UMs were shown to be expressed and function also in skeletal muscle, including myosin I isoforms, myosin VA, myosin VI, myosins XVIIIA and XVIIIB [^1,2^]. Myosin VI (MVI) is a unique UM, which, unlike all other known myosins, moves backwards, i.e., towards the minus end of the actin filament. MVI participates in a variety of intracellular processes such as vesicular membrane traffic, cell migration, adhesion, mitosis, gene transcription, etc [^3, 4^]. MVI is ubiquitously expressed in all metazoans, exhibiting a domain organization similar to all other myosins [^3^]. Its ∼140-kDa heavy chain contains an N-terminal motor domain (with the actin- and ATP-binding sites), a neck, which binds two calmodulin molecules, and a tail domain with its C-terminal part involved in cargo binding [^5^]. It was also shown that a point mutation within *MYO6* is associated with left ventricular hypertrophy [^6^]. Our previous findings provided new information indicating that MVI is an important player in the regulation of heart and skeletal muscle functioning [^2,7^]. We demonstrated that loss of MVI in mice resulted in heart enlargement followed by age-progressing left ventricular remodeling and systolic dysfunction [^7^]. Also, we were the first to show that in myoblasts the mechanisms controlling cytoskeleton organization, myoblast adhesion and fusion are dysregulated in the absence of MVI [^8^].

Interestingly, this molecular motor was shown to be involved in the assembly of F-actin cages to encapsulate Parkin-positive damaged mitochondria and prevention of their refusion with neighboring populations in human embryonic kidney cells (HEK293 cell line). Loss of MVI leads to a mitophagy defect with an accumulation of mitophagosomes and downstream mitochondrial dysfunction [^9^]. These results suggest that MVI is involved in the functioning of mitochondria, the organelles which are an important source of energy for maintaining normal skeletal muscle metabolism. Previously, we demonstrated that lack of MVI impairs cAMP/PKA signaling pathway in skeletal muscle, known to be involved in muscle development and metabolism [^2,10^]. Importantly, it is well-established that the cAMP signaling cascade is interconnected with mitochondrial dynamics and it has been also proposed to be involved in the regulation of oxidative phosphorylation (OXPHOS) [^11,12^]. The abovementioned information encouraged us to undertake the study on the role of MVI in skeletal muscle metabolism. Herein, we have demonstrated for the first time that muscles of MVI-knockout (MVI-KO) mice exhibit increased levels of phospho-AMPK kinase and downregulation of mTOR kinase pathway. The changes in the pivotal metabolic pathways are accompanied by activation of lipolysis and a shift from glycolytic to oxidative fiber types, possibly as a compensatory mechanism to meet the energy demands of MVI-KO muscles. The impairment of mitochondrial integrity and a decrease of ATP levels confirm the compromised mitochondrial status of myogenic cells in the absence of MVI, explaining the activation of AMPK and the downregulation of the mTOR pathway. These findings underscore the important role of MVI in regulation of muscle metabolism.

## 2. Material and Methods

### 2.1. Animals

*Snell’s waltzer* mice (*SV*, C57BL/6J background) were used in the study. These mice do not synthesize MVI due to a 130-bp deletion which results in the introduction of a premature stop codon in the neck region of MVI and therefore serve as a natural MVI knockouts. Each experiment was performed at least three times using a pair of control (heterozygous, WT) and mutant (MVI-KO) mice from the same litter. A sample size was determined from the previous research on the preliminary studies, which provided the magnitude of the difference or relationship being studied. Considerations such as the availability of subjects and practical limitations were taken into account. The mice were bred and housed under pathogen-free conditions in the animal facility of the Nencki Institute. All mice were on the standard rodent chow diet. The decision to use male mice in current study based on similar body weight differences observed between sexes in preliminary data (Supplementary Figure 1), allows for controlled and reproducible results while eliminating confounding variables associated with hormonal fluctuations, estrus cycles, or potential sex-specific responses. Moreover, our previous studies were conducted on male SV mice. Experiments were performed on gastrocnemius (GM), soleus (SOL), extensor digitorum longus (EDL), and tibialis anterior (TA) hindlimb muscles of WT and MVI-KO mice at different age: 3- and 12-months-old (3m and 12m, respectively, male). Experiments were also done on newborn (P0) mice and in this case examination of whole hindlimb muscles of both males and females was performed. Animal housing and euthanasia procedures were performed in compliance with the European Communities Council directives adopted by the Polish Parliament (Act of January 15, 2015 on the use of animals in scientific investigations). To work with SV mice, which are natural knockouts, did not require any permission from a local ethics committe but only approvals from the Director of the Nencki Institute of Experimental Biology; the following approvals were granted: 414W/2021/IBD, 155/2020/IBD and 410W/2021/IBD.

### 2.2. Cell culture

C2C12 Scrambled (Scr) and MVI-KD mouse myoblasts were cultured in Dulbecco’s Modified Eagle’s Medium (DMEM, Gibco, 31966021) with 4.5 g/l glucose. The culture medium was supplemented with 10% heat-inactivated fetal bovine serum (FBS, Gibco, 10082147), antibiotics (1000 UI/ml penicillin and streptomycin 1000 UI/ml, Gibco, 15140122) with the additional presence of 0.4 mg/ml hygromycin B (BioShop, HYG002). The cells were maintained at 37 °C in a humidified atmosphere with 5% CO2.

### 2.3. Myoblasts primary culture

Cell culture enriched with myoblasts, referred to here as primary myoblast culture, were sourced from the hindlimb muscles (gastrocnemius, tibialis anterior, soleus, and extensor digitorum longus muscles) of three-month-old *Snell’s waltzer* mice. The muscles were dissected, and the dissociation process was carried out using 0.2% collagenase (Sigma Aldrich, C0130) as described in [^8^]. Once the cells reached confluence, they were seeded in an appropriate density on tissue culture plastics (depending on the experimental requirements) coated with 10% Matrigel (Corning, CLS354234) and used for further study.

### 2.4. siRNA knockdown of MVI

MVI-knockdown (MVI-KD) stable cell line was generated based on the pSilencer 2.1-U6 hygro vector system (Ambion Inc., USA) essentially as described by [^13^]. MVI protein level was assessed by Western blot analysis.

### 2.5. Analysis of energy metabolism in myoblasts

The oxygen consumption rate (OCR) and ATP production rate were measured using an XF96 analyzer (Seahorse Bioscience) and Agilent Seahorse XF Cell Mito Stress Test Kit (Agilent, 103015-100) which allows real-time determination of oxygen (O_2_) concentration and glycolytic extra-cellular acidification rates (ECAR). C2C12 MVI-KD and Scr myoblasts as well as MVI-KO and WT primary myoblasts were seeded at 10,000 cells/well in 80 µl of appropriate medium in Seahorse XF96 cell culture microplates (Agilent Technologies, 103792-100) and incubated for 24 h at 37°C and 5% CO2. The OCR and ECAR measurements were taken according to the manufacturer’s instructions simultaneously under basal conditions and after the addition of mitochondria uncouplers. After measurements of OCR and ECAR, obtained data were normalized to the cell density estimated as absorbance units (AU, directly proportional to the cell number) using CellTiter 96 AQueous Non-Radioactive Cell Proliferation Assay (Promega Corporation, G5421).

### 2.6. Growth response of myoblasts in a galactose medium

C2C12 MVI-KD and Scr myoblasts were seeded in 96-well plates at a density of 1000 cells/well in an appropriate medium (described in 2.2. Cell culture). After 24 h, cell medium was removed and replaced by a galactose medium composed of DMEM without D(+)-glucose (Gibco, 11966025) and supplemented with 10 mM D(+)-galactose (Sigma Aldrich, G5388) as the sole source of carbohydrate, 2 mM glutamine (Gibco, 25030081), 1 mM sodium pyruvate (Gibco, 11360070), 10% FBS (Gibco, 10082147), and 1% penicillin/streptomycin (Gibco, 15140122). Growth curves were obtained using the Incucyte® SX5 (live-cell analysis system) HD live-cell imaging system (Sartorius), which visualized cells in phase contrast every 3 h and analyzed with the IncuCyte 2023A software (Sartorius).

### 2.7. Measurement of intracellular ATP level

Isolated muscles were homogenized in 10 volumes of 0.1 M phosphate buffer pH 7.75. Myoblasts were initially placed in 35-mm cell culture dishes (Sarstedt, 83.3900) at 40,000 cells/plate and grew them until cells achieved 90% confluence. Next, cells were lyzed in 300 µl of thd phosphate buffer in a room temperature. ATP concentration was measured using the ENLITEN® ATP Assay System (Promega Corporation, FF2000) according to the manufacturer’s instructions. Luminescence was recorded with the use of a microplate reader (Infinite M1000pro, Tecan). Data obtained from primary myoblasts were normalized to the protein concentration, determined by the BCA Protein Assay Kit (Thermo Scientific, 23227).

### 2.8. Measurement of mitochondria integrity using flow cytometry

Primary myoblasts were stained using MitoTracker^TM^ Red CMXRos dye (Invitrogen, M46752) at 0.5 µM final concentration according to the manufacturer’s instructions. Flow cytometry was performed on the Guava® easyCyte™ 8HT flow cytometer from EMD Millipore. Data were collected for 10,000 myoblasts each time. Histograms were created using InCyte software (Millipore).

### 2.9. Real-time polymerase chain reaction (RT-qPCR)

RNA was isolated from examined muscles and RT-qPCR was performed as described in [^7^]. The primers for qPCR were designed using primer-BLAST: for *Myo6* 5′-GATCTGTCCCAGCAGGAAGC-3′ (forward) and 5′-TTATCCACTATCTCCCGGCG-3′ (reversed), for *B2m* 5′-CATGGCTCGCTCGGTGACC-3′ (forward) and 5′-AATGTGAGGCGGGTGGAACTG-3′ (reversed), for *Prkaa2* 5′-CGGCAAAGTGAAGATTGGAGAA-3′ (forward) and 5′-TCCAACAACATCTAAACTGCGAAT-3′ (reversed).

### 2.10. Western blot analysis

The muscles were homogenized as described in [^7^]. For estimation of MVI level, 1 M phosphate buffer pH 7.2 supplemented with 1 mM phenylmethanesulfonyl fluoride (Serva, 32395) was used for tissue homogenization. Cells were lysed in an ice-cold RIPA buffer containing 50 mM Tris-HCl pH 7.5, 150 mM NaCl, 0.5% sodium deoxycholate, 0.1% SDS, 1% Nonidet P-40, 50 mM sodium fluoride, and 1 mM PMSF supplemented with protease and phosphatase inhibitors. Muscle homogenates or cell lysates (10–30 μg of protein/well) were separated using 8, 10, or 12% polyacrylamide SDS-gels and then transferred to a nitrocellulose membrane (Amersham, 10600002) followed by immunoblotting as described in [^8^].

The following primary antibodies were used: against GAPDH (glyceraldehyde-3-phosphate dehydrogenase, Merck, MAB374, 1:50000); β-tubulin (Abcam, AB21058, 1:10000); MVI (Proteus Biosciences, 25-6791, 1:1000); AMPK (AMP-activated protein kinase, Cell Signaling, 2532, 1:1000); p-AMPK (Thr172) (Cell Signaling, 2535, 1:1000); mTOR (mammalian target of rapamycin, Cell Signaling, 5536, 1:500); p-mTOR (Ser2448) (Cell Signaling, 5536, 1:500); p70S6K (ribosomal protein S6 kinase, Cell Signaling, 2708, 1:1000); p-p70S6K (Thr389) (Cell Signaling, 9234, 1:250); S6 (S6 ribosomal protein, Cell Signaling, 2217, 1:5000); pS6 (Ser 235/236), Cell Signaling, 4858, 1:5000); 4E-BP1 (Cell Signaling, 9452, 1:500); p-4E-BP1 (eukaryotic translation initiation factor 4E (eIF4E)-binding protein 1, Ser65) (Cell Signaling, 9451, 1:500); ATGL (adipose triglyceride lipase, Cell Signaling, 2138, 1:500); HSL (hormone-sensitive lipase, Cell Signaling, 4107, 1:500); p-HSL (Ser565) (Cell Signaling, 4137, 1:500); p-HSL (Ser563) (Cell Signaling, 4139,1:500); PLIN1 (perilipin 1, Abcam, ab60269, 1:250). The following secondary antibodies were used: HRP-conjugated anti-mouse IgG (Millipore, AP308P); HRP-conjugated anti-rabbit IgG (Millipore, AP307P); HRP-conjugated anti-goat IgG (Millipore, AP186P).

### 2.11. Muscle fiber type composition analysis

Muscle tissues embedded in Optimal Cutting Temperature compound (Thermo Fisher Scientific; 6769006) were frozen in liquid nitrogen-cooled 2-methylbutane (VWR Chemicals, 103616V). Cryosections (10 µm) were obtained on Leica CM 1950 cryostat. Immunohistochemical detection of myosin heavy chain (MyHC) was performed as previously described [^14^]. Primary and secondary antibodies used are listed in Table S3. Images were acquired with a fluorescent Olympus VS110 scanning microscope equipped with an RGB camera and U-PLAN 20×/0.75 dry objective. Fiber type distribution was quantified using ImageJ [^15^].

### 2.12. Immunolocalization study

Cells or muscle fibers were fixed in 4% formaldehyde in PBS pH 7.4 for 30 min at room temperature, washed with PBS, permeabilized with 0.3% Triton X-100, and blocked in 2% goat serum. Samples were then incubated overnight at 4°C with primary antibodies cocktail against MyHC I (Developmental Studies Hybridoma Bank; BA-F8), IIa (DSHB; SC-71), IIb (DSHB; BF-F3), and dystrophin (Abcam; AB7164) to stain different muscle fiber types. After washing step a secondary antibodies mix consisting of Alexa Fluor 488 Goat Anti-Mouse IgG1 (Thermo Scientific; A-21121), Alexa Fluor 555 Goat Anti-Mouse IgM (Thermo Scientific; A-21426), and Alexa Fluor 633 Goat Anti-Mouse IgG2b (Thermo Scientific; A-21146) was applied. Vectashield mounting medium (Vector Laboratories, H-2000) was used to mount the slides. Images were collected with Zeiss LSM780, Inverted Axio Observer Z.1 equipped with 63×/1.4 Oil Plan Apochromat DIC objective and processed using the Zen Blue 2.1 software (Carl Zeiss Microscopy). A quantitative assessment of fluorophore co-localization in confocal optical sections was performed using Pearson’s correlation coefficient, which is a well-defined and commonly accepted tool for describing the extent of overlap between image pairs [^16^]. The value of this coefficient ranges from −1 to 1, with a value of −1 representing a total lack of overlap between pixels from the images, value of 1 indicating perfect pixel overlapping, and a value 0 indicating no correlation. Areas were randomly selected and the fluorescence image profile was obtained using Fiji distribution of ImageJ 1.52a software.

### 2.13. Skeletal muscle triacylglycerol measurement

Muscle tissues were homogenized in 1ml of 5% IGEPAL CA-630 (Merck, I3021). The samples were then heated at 90°C for 5 minutes, cooled to room temperature and this step was repeated one more time. Next, the samples were centrifuged for 2 min at 14 000 x g followed by 10 times dilutions. Triglyceride content was quantified utilizing a Triglyceride Quantification Colorimetric Kit (Sigma-Aldrich; MAK266) following the manufacturer’s instructions.

### 2.14. Statistics

All experiments were done at least three times in two or three technical replicates. Results were expressed as means ± SD (standard deviation). The number of animals was indicated in the graphs (total number of mice used for study - 115). If the data were normally distributed we performed parametric two-tailed Student’s *t*-test or One-way ANOVA in GraphPad Prism 8.4.3 software (San Diego, CA, USA). Data that were non-normally distributed were analyzed with a non-parametric Mann-Whitney test to determine significance. Statistical significance was defined as \**p*<0.05, \*\**p*<0.01, *** or ^&&&^*p*<0.001,****, ^&&&&^ or ^####^*p*<0.0001.

## 3. Results

### 3.1. MVI expression in different skeletal muscles throughout muscle development

Our previous findings showed that MVI is expressed in rodent and human skeletal muscles, C2C12 murine myoblasts, primary myoblasts (isolated from WT mice), and both nascent and mature myotubes [^8,17,18^]. Its synthesis gradually decreases during differentiation [^8,13^]. To gain deeper insight into MVI expression in skeletal muscles throughout the animal lifespan, we performed RT-qPCR (to measure Myo6 mRNA levels) and Western blots (to estimate MVI protein level) in hindlimb muscles at various developmental mouse stages, namely in P0, 3m and 12m mice.

Analysis of the mRNA level revealed that the highest expression of *Myo6* was observed in P0 muscles as illustrated in Figure 1A, followed by a decrease in 3m and 12m. This is in line with our recent studies on murine hearts [^7^]. Similar results were obtained on the protein level (Figure 1B and C). Notably, the SOL muscle, characterized by the highest percentage of slow-oxidative fibers, exhibited the highest MVI protein level (Figure 1B and C). Conversely, the lowest expression of *Myo6* gene and MVI content was observed in the 12m TA muscle.

**Fig 1.**
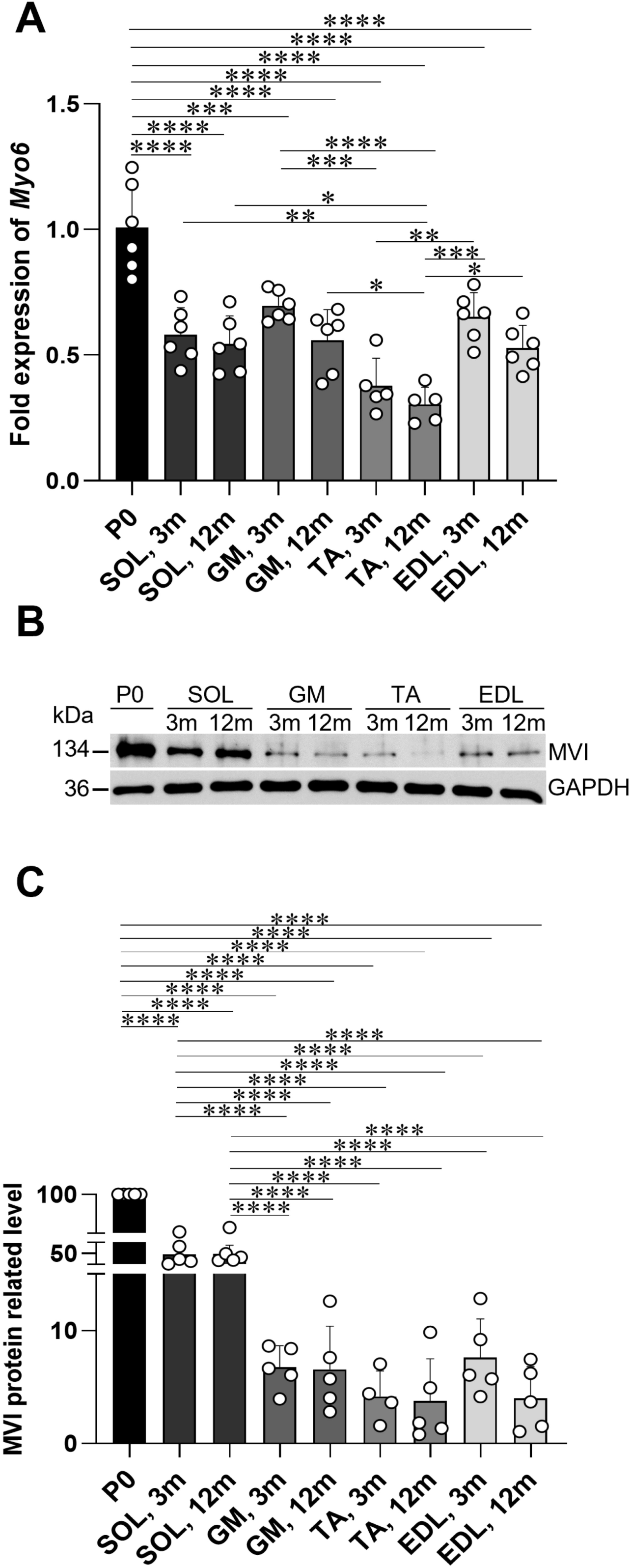
MVI expression in hindlimb muscles. (A) Relative expression of mRNA-*Myo6* in gastrocnemius (GM), soleus (SOL), extensor digitorum longus (EDL), and tibialis anterior (TA) muscle of different age: newborn (P0), adult 3-months-old (3m) and 12-months-old (12m) WT mice. mRNA-*B2m* (β-2 microglobulin) was used for normalization. Statistical significance was analyzed with One-way ANOVA test, *p < 0.05, **p < 0.01, ***p < 0.0001, ****p < 0.0001 (P0 served as 1). (B) Representative Western blot analysis and (C) corresponding densitometry analyses of MVI protein level. GAPDH was used as a protein loading control. Statistical significance was analyzed with One-way ANOVA test, ****p < 0.0001 (P0 WT served as 1).

### 3.2. Glycolytic-to-oxidative fiber-type switch in skeletal muscles under the absence of MVI

Previously observed by us alterations in the cAMP/PKA pathway contributed to the significant decrease in muscle fiber size observed in MVI-KO mice [^2^]. To investigate the fiber-type composition of muscles under the lack of MVI, we performed muscle fiber typing on the four examined muscles with the use of the myosin heavy chain (MyHC) immunohistochemistry. The analysis revealed a consistent trend of transitioning from glycolytic to oxidative fiber types in all examined MVI-KO muscles (Figure 2A–D, Supplementary Figure 2A, B). Specifically, in SOL muscles isolated from 3m MVI-KO animals we observed a tendency towards an increase in type IIa (fast-oxidative) fibers. This glycolytic-to-oxidative transition was even more pronounced in 12m MVI-KO SOL muscles (Figure 2A, B), reflected in a significant increase in the percentage of type I fibers (slow-oxidative). EDL muscle, known for its abundance of type IIb/x fibers (fast glycolytic), exhibited an elevation in the presence of type IIa fibers in both 3m and 12m mice lacking MVI, compared to WT animals (Figure 2C, D). A similar trend was observed in the muscles with mixed fiber types, i.e. TA and GM (Supplementary Figure 2A, B), although the observed alterations were relatively minor in TA muscle, where the MVI expression is the lowest.

**Fig 2.**
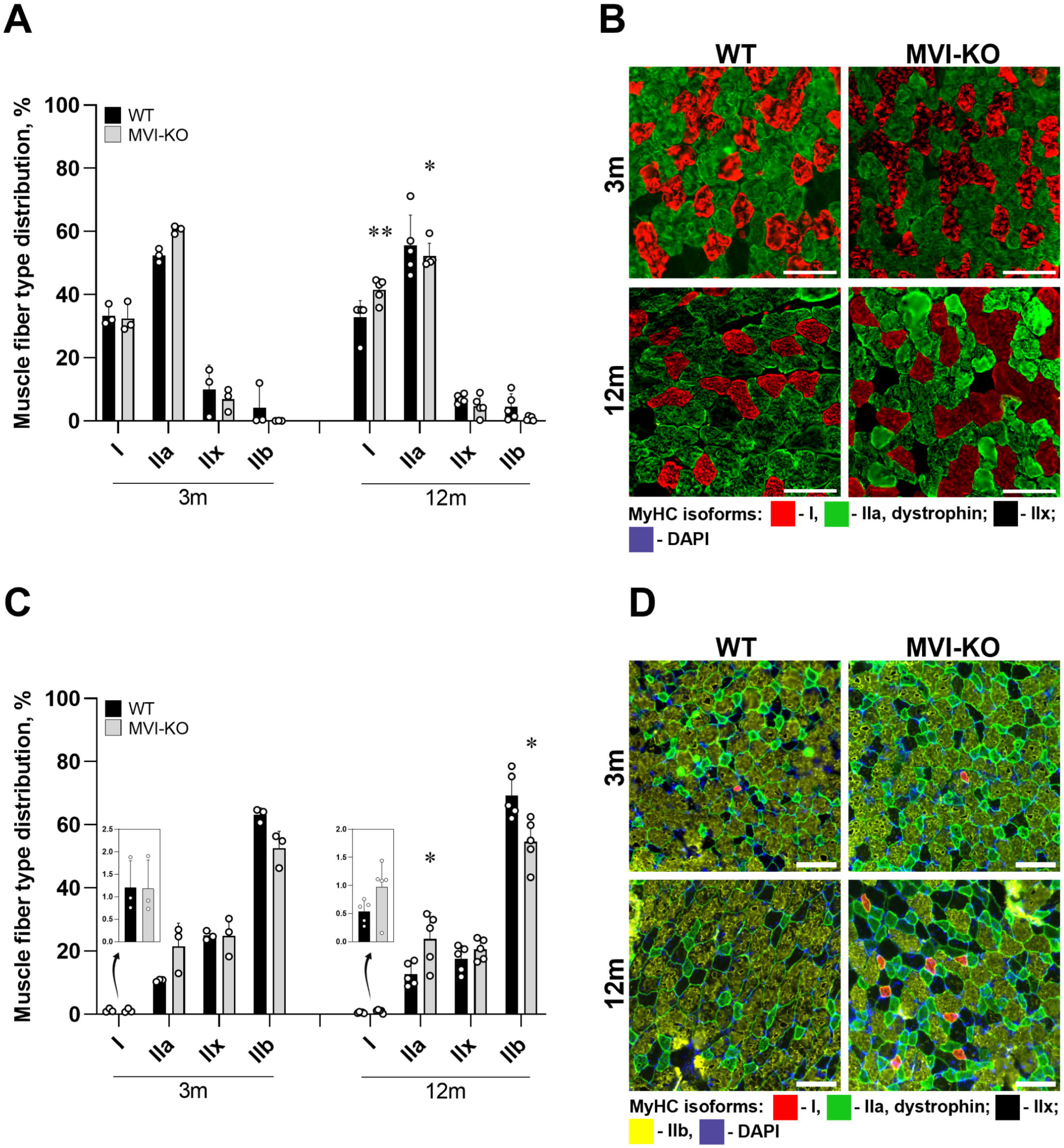
Glycolytic-to-oxidative fiber-type switch in MVI-KO muscles. (A) Fiber typing *via* myosin heavy chain (MyHC) immunohistochemistry in MVI-KO and WT Soleus (SOL) muscle isolated from 3- and 12-months-old animals. (B) Representative microphotographs of immunofluorescence staining of MVI-KO and WT SOL muscle. (C) Fiber typing *via* MyHC immunohistochemistry in MVI-KO and WT extensor digitorum longus (EDL) muscle. Inserts, Graphs representing the percentage of type I muscle fibers, pointed by arrow. (D) Representative microphotographs of immunofluorescence staining of MVI-KO and WT EDL muscle. Bars, 100 µm. Statistical significance was analyzed with t-test, *p < 0.05, **p < 0.01, vs. WT.

### 3.3. MVI and TOM20 localization in myogenic cells and muscle fibers

The observed glycolytic-to-oxidative transition of the examined MVI-KO muscles, particularly in SOL muscle, indicates a possible involvement of MVI in muscle metabolism and mitochondria function. To address this issue, we performed immunofluorescence staining for MVI and TOM20 (a marker of mitochondrial mass due to its constitutive expression in the outer mitochondrial membrane) in myogenic cells and muscle fibers. As shown in Figure 3, double staining for MVI (green) and TOM20 (red) revealed a partial co-localization of both proteins in myogenic cells (myoblasts, myocytes, and mytotubes (Figure 3A–C) as well as in muscle fibers (isolated from 3m and 12m SOL WT mice (Figure 3D, E)). This was confirmed by a Pearson co-localization analysis as the value of Pearson’s coefficient (r_p_) for the images exceeded 0.5 in all examined cases.

**Fig 3.**
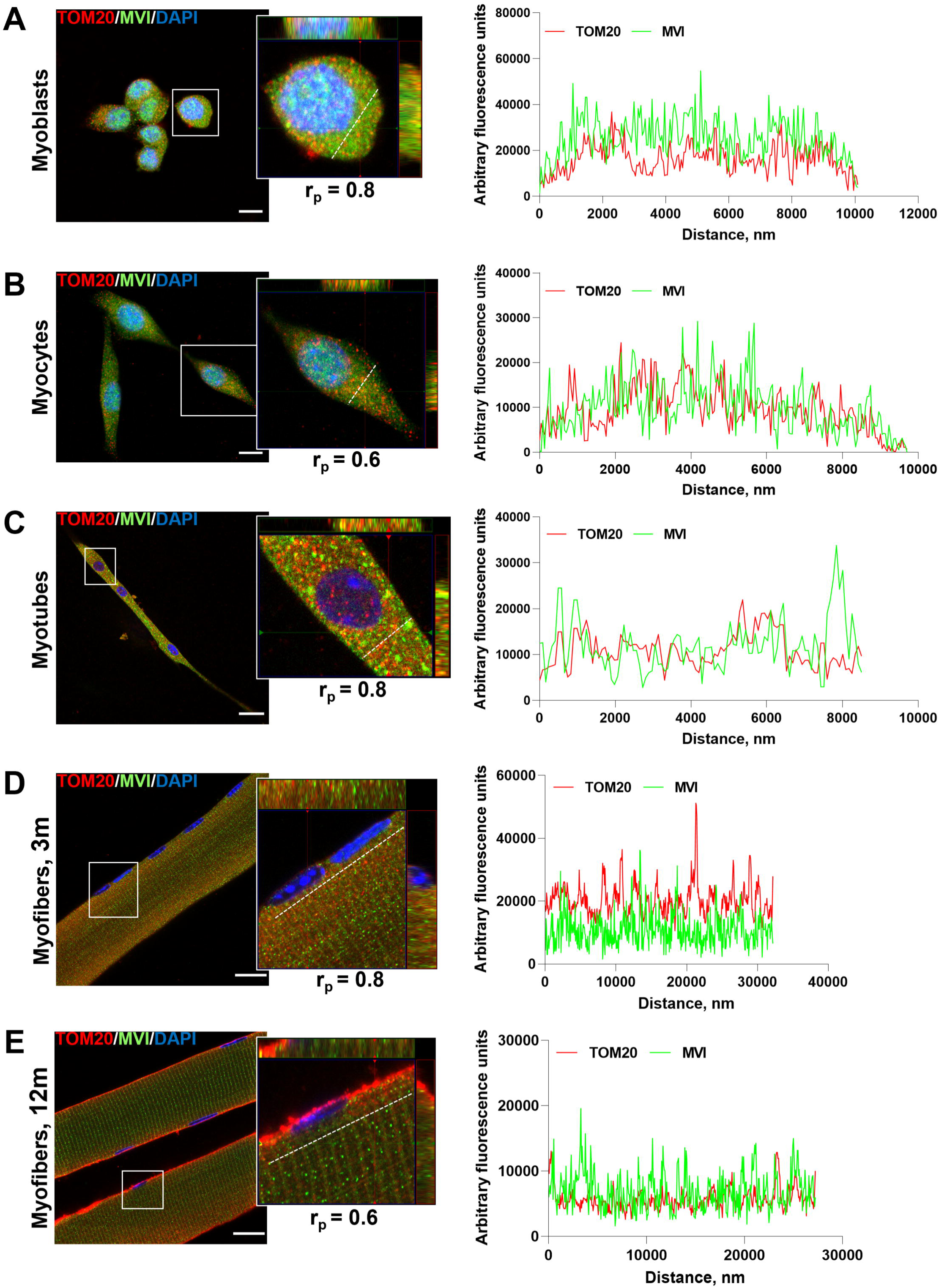
MVI and TOM20 co-localization in myogenic cells and muscle fibers. Distribution (left panel) and fluorescence image profile (right panel) of MVI and TOM20 in (A) myoblasts; (B) myocytes; (C) myotubes; (D) myofibers isolated from SOL muscle of 3-months-old mice; (E) myofibers isolated from SOL muscle of 12-months-old mice. Dashed line indicates the site where fluorescence profile was measured. Graphs represent fluorescence intensity profiles calculated on corresponding images obtained from samples co-immunostained for MVI and TOM20. Bars, 10 μm in A, B, D, E and 20 μm in C. R_p,_ Pearson’s correlation coefficient, calculated in corresponding samples.

### 3.4. Mitochondrial dysfunction in myogenic cells under the lack of MVI

To further investigate a link between MVI and mitochondria, we used the Optimized Seahorse Mito Stress assay in murine myoblast cells of MVI-KD C2C12 line. Basal respiration was considerably lower in MVI depleted cells (knockdown rate was 92%) compared to the control Scr cells (****p<0.0001, Figure 4A, Supplementary Figure 3A). This observation indicates a lower reliance on OXPHOS for energy production in MVI-KD cells. Also, MVI knockdown resulted in a marked reduction of the maximal respiration as compared to Scr cells (****p<0.0001, Figure 4A, Supplementary Figure 3B).

**Fig 4.**
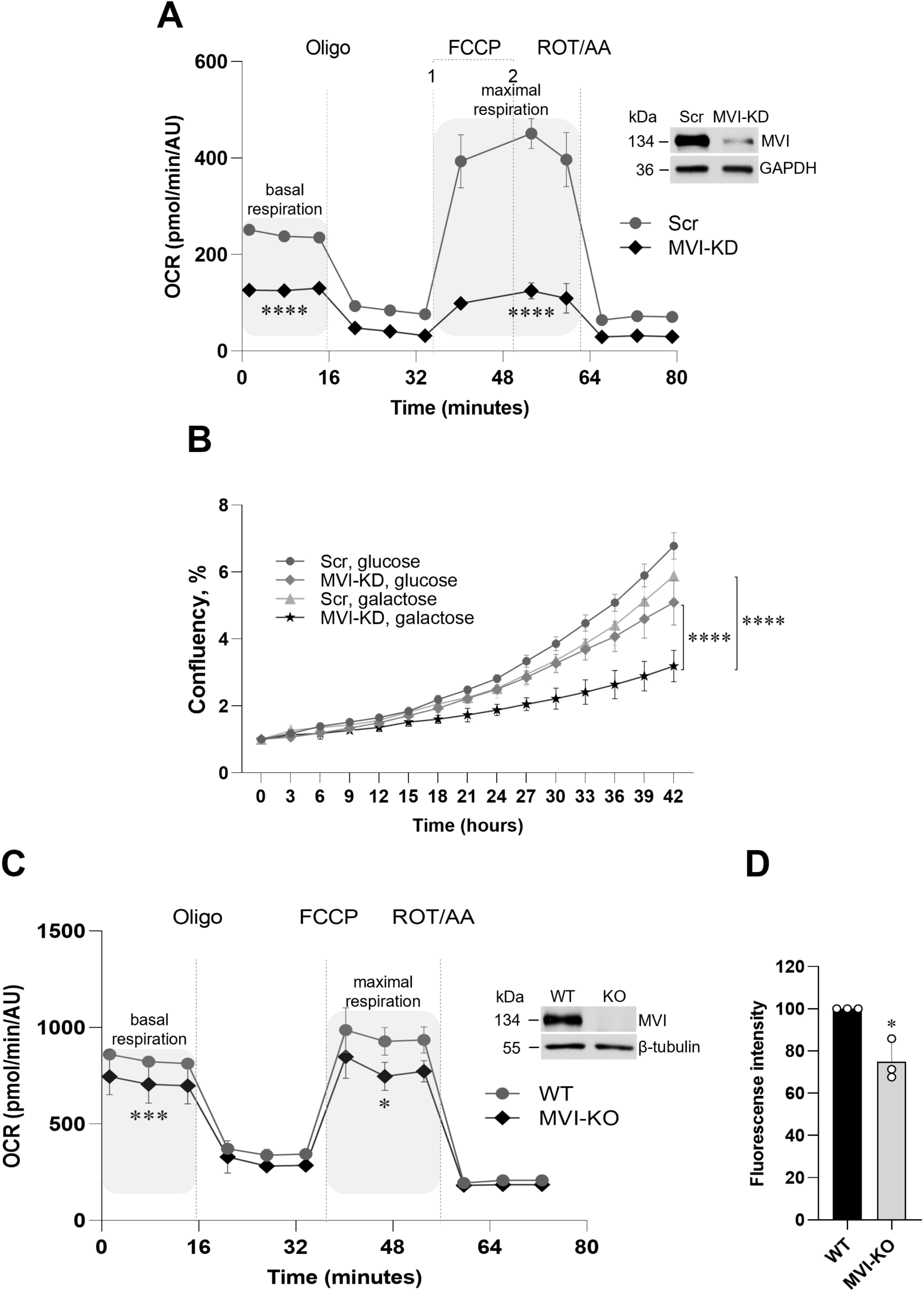
Mitochondria dysfunction in myogenic cells under the lack of MVI. (A) Measurement of the oxygen consumption rate (OCR) in C2C12 murine myoblast cells. Scr, Control; MVI-KD, stable MVI knockdown (See Methods for details). Oligomycin (Oligo, 1 µM), fluoro-carbonyl cyanide phenylhydrazone (FCCP, 1 – 0.5 µM or 2 – 0.75 µM), rotenone (ROT, 1 µM), and antimycin (AA, 5 µM) were injected at the indicated times to determine OCR. The steady state OCR was defined as the baseline OCR before any reagent has been added. Results from these experiments were used to determine the basal respiration (defined as the differences between the OCR of the basal conditions and non-mitochondrial OCR after ROT/AA treatment) and maximal respiration (defined as the maximum OCR after FCCP uncoupling). Data were normalized to the cell density estimated as absorbance value. Insert, immunoblotting for MVI in C2C12 Control and MVI-KD cells. The knockdown rate was 92%. GAPDH served as loading control. (B) Growth of MVI-KD and Scr cells in glucose and galactose medium. (C) Measurement of the oxygen consumption rate (OCR) in primary myoblasts. WT – cells isolated from muscles of heterozygous mice, MVI-KO – cells isolated from muscles of MVI-knockout mice. Oligo 1 µM, FCCP, 0.75 µM, ROT 1 µM, and AA, 5 µM were injected at the indicated times to determine OCR. Data were normalized to the cell density estimated as absorbance value. Insert, immunoblotting for MVI in WT and MVI-KO cells. β-tubulin served as a loading control. (D) Mitochondrial membrane potential was measured using MitoTracker^TM^ Red CMXRos dye by flow cytometry. Data are represented as mean ± SD, n = 3. Statistical significance was analyzed with t-test, *p < 0.05, ***p ≤ 0.001 and ****p ≤ 0.0001 vs. Control.

To further assess mitochondrial function under MVI knockdown, we compelled Scr and MVI-KD C2C12 cells to depend on mitochondrial respiration as their main energy source [^19^]. To achieve this, cells were cultured in a medium with galactose as the exclusive carbon source, instead of glucose. We observed that proliferation of MVI-KD cells in galactose medium exhibited a notable statistically relevant inhibition (****p<0.0001 compared to MVI-KD in glucose medium; **p<0.01 compared to Scr in galactose medium), as evidenced by increased doubling times (Figure 4B), indicating a mitochondrial defect. We also noticed a tendency for slightly higher Scr cell growth rates in glucose compared to galactose, as depicted in Figure 4B. This phenomenon can be explained by the difference in galactose uptake compared to glucose since glucose transporters have a higher affinity for glucose than galactose [^20,21^]. A modest decrease in the growth rate of MVI-KD cells compared to Scr in glucose medium was noticed. A similar phenomenon was described in MVI-KD HeLa and HEK293 cells and explained by cytokinesis defect of these cells [^9,22^].

To examine the impact of the complete MVI loss on mitochondria, we measured OCR in myoblasts derived from MVI-KO mice. Remarkably, cells without MVI also showed a significant reduction in basal and maximal respiration compared to control cells (****p<0.0001, Figure 4C, Supplementary Figure 3C and D), indicating a compromised mitochondrial function.

Moreover, to confirm the mitochondrial dysfunction in MVI-KO myoblasts, we examined the mitochondrial status through MitoTracker Red staining, followed by FACS analysis (Figure 4D). Indeed, a noticeable reduction (from 100% in WT to 75% in MVI-KO) in mitochondrial integrity was observed in MVI-KO myogenic cells.

### 3.5. Loss of MVI affects ATP production

Mitochondrial dysfunction usually results in a subsequent compromise of the bioenergetic balance. To further substantiate a link between mitochondria impairment under the lack of MVI and cellular bioenergetics, the ATP production rate was assessed to monitor mitochondrial respiration and glycolysis of myoblasts (Figure 5). A significant reduction in a total ATP production, encompassing ATP generated from OXPHOS (mitoATP), and glycolysis (glycoATP), was observed in MVI-KD cells (Figure 5A). However, the ratio between mito- and glycoATP was the same in MVI-KD and Scr line. Quantification of ATP production in primary myoblasts isolated from hindlimb muscles of MVI-KO mice (Figure 5B) revealed that primary myoblasts exhibited a greater reliance on OXPHOS for ATP production. More than a twice increase in the ratio of mitoATP for both WT (77%) and MVI-KO (78%) myoblasts was observed compared to C2C12 cells (31% for Scr and 32% for MVI-KD) (Figure 5B), which predominantly utilizes glycolysis as its primary source for ATP synthesis. However, a similar trend was observed concerning the ATP generation ratios. A pronounced decrease in a total ATP production was also observed in MVI-KO myoblasts compared to WT; analogous observation was made for both mito- and glycoATP ratios in myoblasts lacking MVI. These findings were further validated by assessing intracellular ATP levels using the ENLITEN® ATP Assay System (Figure 5C). A noticeable decrease was observed in the overall ATP production in MVI-KO myoblasts compared to WT cells.

**Fig 5.**
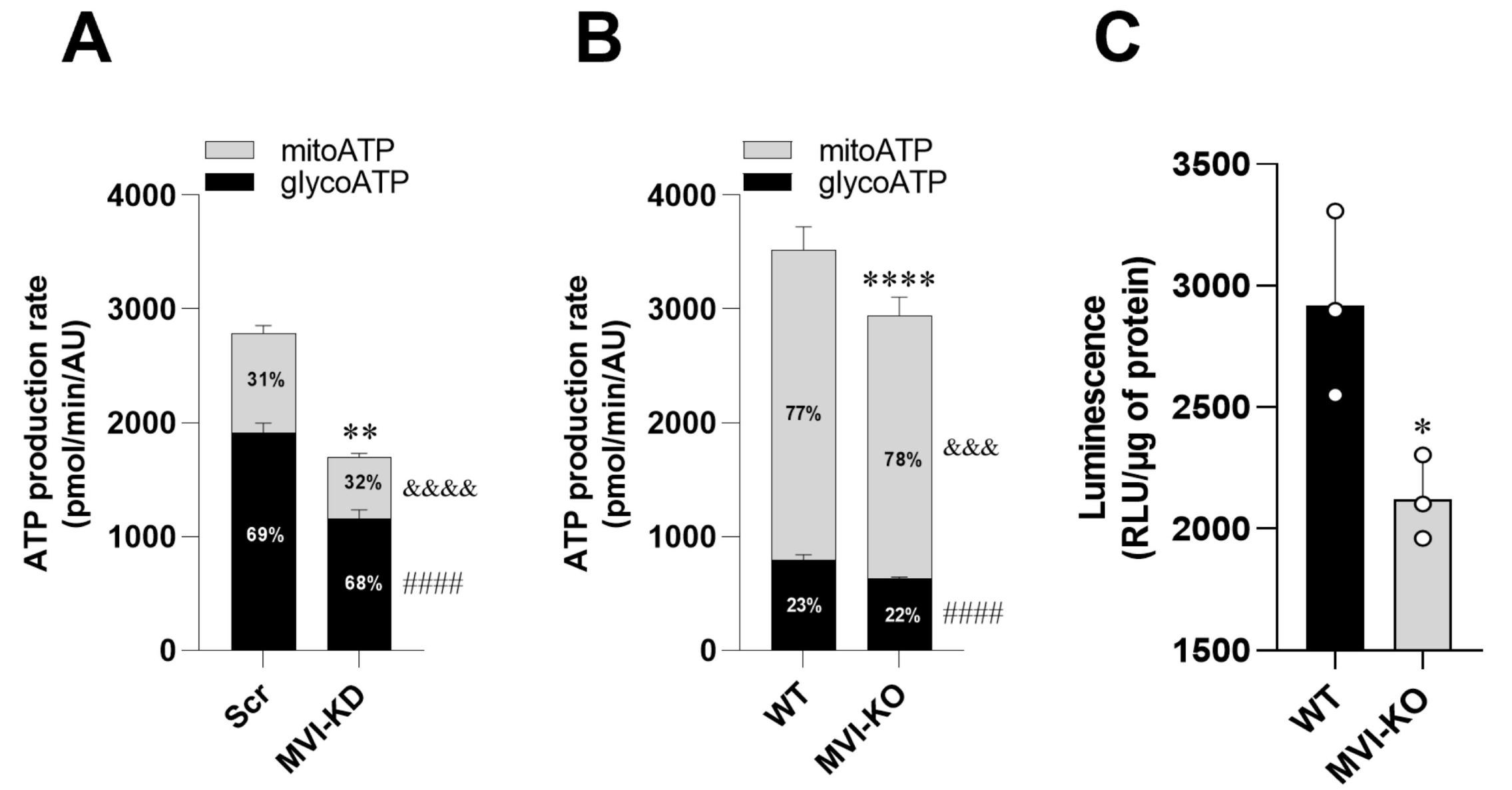
Impairment of ATP production in myogenic cells under the lack of MVI. (A) Real-time ATP production rate in C2C12 murine myoblast cell line. (B) Real-time ATP production rate in primary myoblasts. Data are represented as mean ± SD, N = 3, n (technical replicates) ≥ 8 (C) Measurement of total intracellular ATP level in primary myoblasts. Data are represented as mean ± SD. Statistical significance was analyzed with t-test, *p ≤ 0.05 vs WT, **p ≤ 0.01 vs Scr, ****p ≤ 0.0001 vs. WT, ^&&&^p t-test ≤ 0.001 compared to mitoATP production rate of WT cells, ^&&&&^p t-test ≤ 0.0001 compared to mitoATP production rate of Scr cells, ^####^p t-test ≤ 0.0001 compared to glycoATP production rate of Scr or WT cells.

Next, we assessed ATP levels in muscles from MVI-KO mice (Figure 6A, Supplementary Figure 4). Our findings revealed a 50% reduction in ATP content within MVI-KO SOL muscle from 3m mice (Figure 6A), the muscle type for which we observed the highest MVI protein level in control animals (Figure 1B and C). Furthermore, the ATP level continued to decrease (up to 21%) in SOL muscles isolated from 12m MVI-KO mice when compared to their wild-type counterparts (Figure 6A). The other examined muscles (GM, TA and EDL) exhibited a similar trend (Supplementary Figure 4).

**Fig 6.**
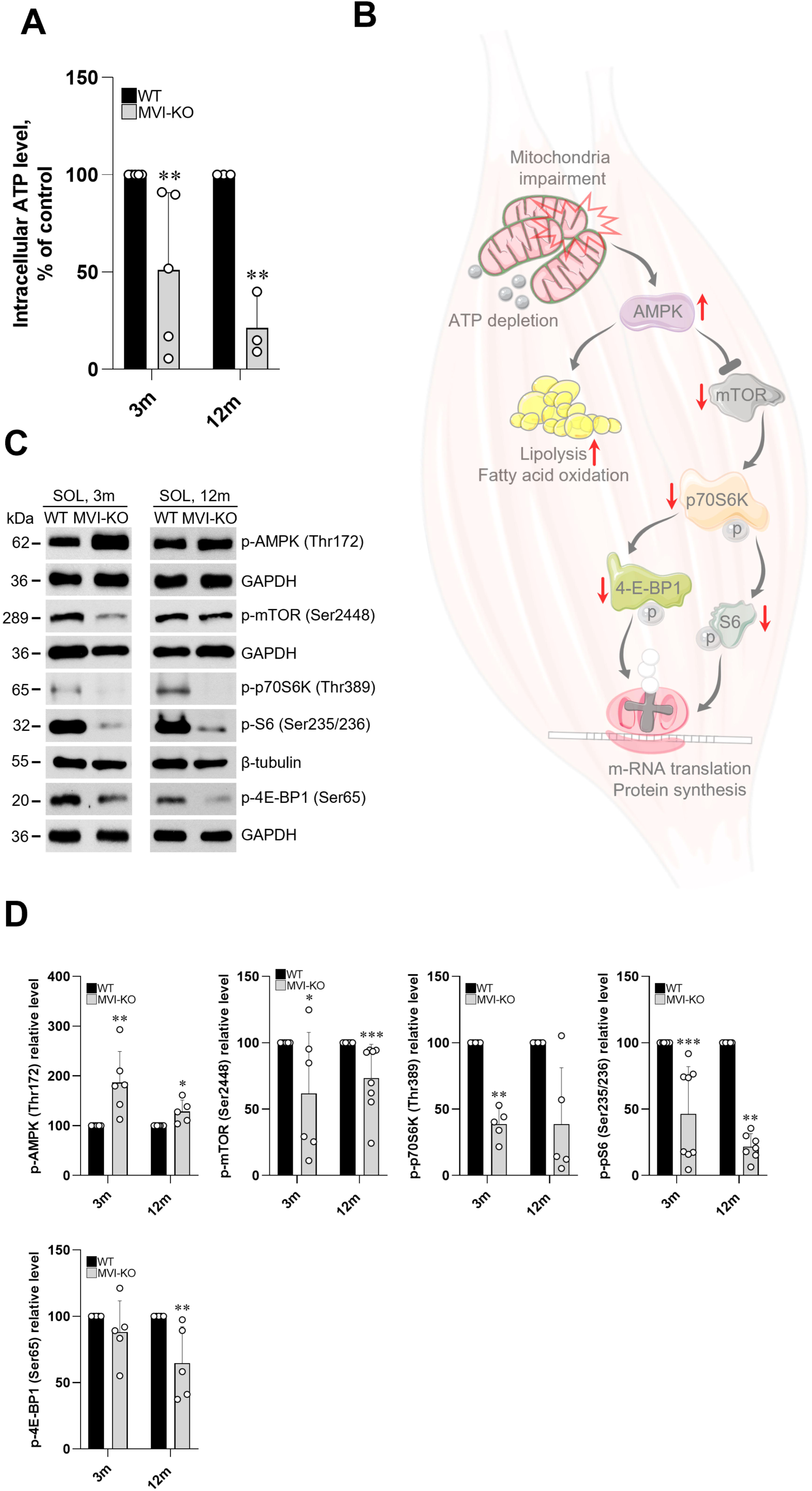
Effect of MVI loss on ATP production and muscle metabolism. (A), Reduction of intracellular ATP level in MVI-KO Soleus (SOL) muscle isolated from 3- and 12-months-old mice. (B) Schematic representation of the AMPK/mTOR signaling pathway. Alterations in mitochondria homeostasis and ATP depletion under the lack of MVI lead to AMPK activation. Activated AMPK, in turn, initiates a set of biological events aimed at restoring cellular metabolic balance including suppression of mTOR, an important regulator of protein translation, induction of fatty acid oxidation, and lipolysis. Red arrows indicate changes in protein levels under the lack of MVI, pointing either to decreased or increased levels. (C) Representative Western blot and (D) densitometry analyses of key proteins involved in AMPK/mTOR pathway in SOL muscle of MVI-KO and WT animals. GAPDH or β-tubulin served as loading controls. Statistical significance was analyzed with t-test, *p < 0.05, **p < 0.01, ***p < 0.001 vs. WT. Illustration was created using smart.servier.com.

Additionally, we observed an increased level of active (phosphorylated) form of AMP-activated protein kinase (AMPK), an intracellular sensor for ATP consumption and critical regulator of skeletal muscle metabolism, in MVI-KO mice in all examined time points (Figure 6B–D) [^23,24^]. This observation was not associated with the increased expression of AMPK catalytic unit (*Prkaa2* gene, Supplementary figure 5). Also, a subsequent reduction in phosphorylated form of mTOR resulted in a decrease in phosphorylation of its downstream targets, including the active forms of phospho-S6 ribosomal protein (pS6), phospho-ribosomal protein S6 kinase (p70S6K) and eukaryotic translation initiation factor 4E (eIF4E)-binding protein 1 (4E-BP1) indicating alterations in metabolism and downregulation of protein synthesis (Figure 6B–D).

It is known that MVI-KO mice display the circling and hyperactivity phenotype [^25^]. To investigate whether the alterations in ATP production and metabolism in adult mice may be attributed solely to the absence of MVI rather than to their hyperactive phenotype, we performed analogous studies on the muscles of newborns, which also revealed a significant decrease in ATP content in MVI-KO P0 mice (Figure 7A), similarly to adult animals. Additionally, we observed an increase in AMPK phosphorylation, along with a reduction in the mTOR downstream target p-S6 in MVI-KO mice compared to WT (Figure 7B, D). Furthermore, examination of the muscle fiber type composition in newborns revealed a similar trend to that seen in adult muscles, i.e. the muscle fiber profile of MVI-KO hindlimb muscles underwent a shift, displaying an increased abundance of fiber type I accompanied by a decrease in proportion of IIa, IIb, and IIx fibers, indicative of an increase in oxidative metabolism (Figure 7C).

**Fig 7.**
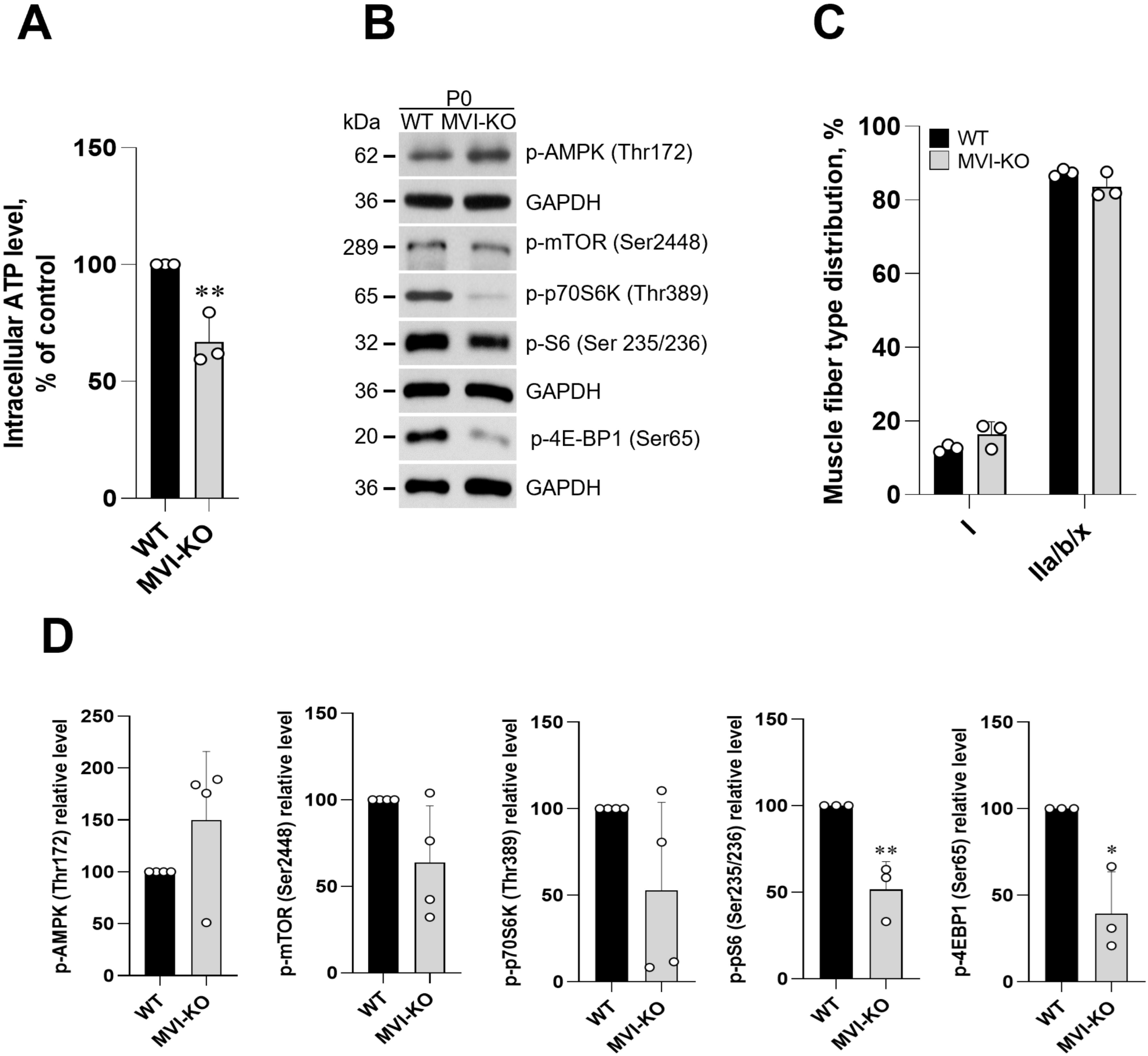
Effect of MVI loss on ATP production and muscle metabolism in hindlimb muscles isolated from newborn (P0) mice. (A) Decrease of intracellular ATP level in MVI-KO P0 hindlimb muscles. (B) Representative Western blot analysis of key proteins involved in AMPK/mTOR pathway in hindlimb muscles isolated from MVI-KO and WT animals. GAPDH served as a loading control. (C) Fiber typing via myosin heavy chain (MyHC) immunohistochemistry in MVI-KO and WT hindlimb muscles isolated from P0 mice. (D) Densitometric analysis of Western blot represented in (B). Statistical significance was analyzed with t-test, *p < 0.05, **p < 0.01 vs. WT.

### 3.6. Lack of MVI resulted in altered lipid metabolism in muscle

During stress conditions, AMPK is triggered to impede processes associated with energy consumption, such as fatty acid and steroid synthesis, while simultaneously activating energy metabolism pathways, including lipolysis and/or fatty acid oxidation (Figure 8A) [^23^]. In light of this, we hypothesized that lipid metabolism in MVI-KO muscles could be altered. Western blot analysis of the SOL muscle was conducted to assess the level and/or activity of lipolysis-related proteins activated through the AMPK pathways in MVI-KO mice compared to their WT counterparts (Figure 8B and C). We observed a significant upregulation of adipose triglyceride lipase (ATGL, playing a crucial role in initiating triacylglycerols (TAG) hydrolysis in skeletal muscle). Also, we showed an increase in the level of hormone-sensitive lipase (HSL) phosphorylated on serine 565 (Ser565) and perilipin 1 (PLIN1), while the level of HSL phosphorylated on serine 563 (Ser563) was decreased. Moreover, we found that a concentration of TAG was significantly reduced in SOL muscle of MVI-KO isolated from both 3m and 12m animals compared to the control, which is the result of accelerated lipolysis (Figure 8D).

**Fig 8.**
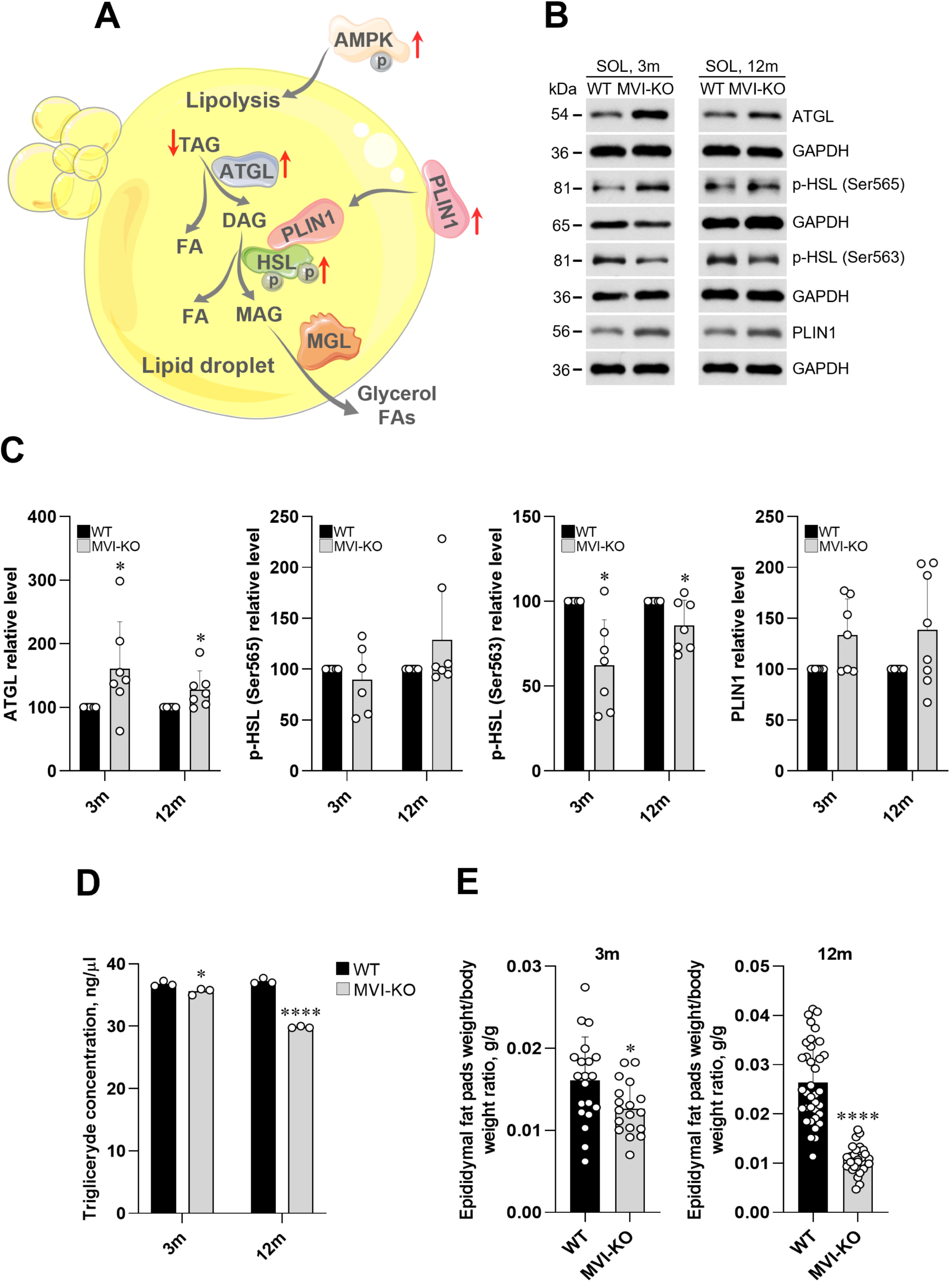
Alterations in muscle lipid metabolism under the lack of MVI. (A) Schematic representation of lipolysis, the hydrolysis of triacylglycerol (TAG) to release fatty acids (FAs). Activation of lipases (hormone-sensitive lipase (HSL), adipose triglyceride lipase (ATGL), and monoacylglycerol lipase (MGL) is crucial for the initiation of this process. ATGL initiates lipolysis by cleaving the first FA from TG and then HSL and MGL act on diacylglycerol (DAG), also releasing FA and glycerol. Activation of AMPK under the lack of MVI resulted in upregulation of ATGL and phosphorylation of HSL on serine 565 residue. Increased level of perilipin 1 (PLIN1) fasilitates lipolisis by increasing the ability of HSL to associate with the lipid droplet surface. Red arrows indicate changes in protein levels, pointing either to decreased or increased levels. (B) Representative Western blot and (C) densitometric analysis of key proteins involved in lipolysis in SOL muscle isolated from 3- and 12-months-old mice. GAPDH served as loading control. (D) Reduced triacylglycerol concentration in MVI-KO Sol muscle of 3- and 12-months-old animals. (E) Decreased epididymal fat pads/body weight ratio in MVI-KO 3- and 12-months-old mice. Statistical significance was analyzed with t-test, *p < 0.05, ****p < 0.0001 vs. WT. Illustration was created using smart.servier.com.

Interestingly, a visual examination of MVI-KO mice revealed that aside from their diminished weight, there was a noticeable decrease in the adipose tissue mass. Specifically, the weight of epididymal fat pads obtained from both 3m and 12m mice was considerably reduced, as illustrated in Figure 8E (normalized to the total body weight), providing additional confirmation of dysregulated metabolic processes in MVI-KO mice.

## 4. Discussion

Herein, we demonstrated for the first time important role of MVI in regulation of muscle metabolism. Expression of MVI in skeletal muscles depends on the muscle type and gradually decreases with age. MVI loss leads to a shift towards oxidative respiration accompanied by a dysregulation of mitochondria function, an increase of the presence of slow-twitch fibers, a decrease in ATP production, a downregulation of AMPK and mTOR pathways as well as an activation of lipolytic enzymes.

SOL, a slow-twitch muscle, contained the highest MVI protein level, while TA, a fast-twitch muscle was characterized by the lowest MVI protein level. Interestingly, SOL muscle was the one to be mostly affected by the MVI loss. Also, it was shown that MVI interacts with mitochondria and is selectively recruited to damaged mitochondria in nonmuscle cells via interaction with Parkin [^9^]. Of note, unconventional myosin XIX (myo19) was shown to be involved in mitochondria functions and network organization [^26,27^]. Herein, we demonstrate that either loss or depletion of MVI compromises mitochondrial function in myogenic cells and affects cell growth in galactose medium, which further confirms mitochondria dysfunction impairing energy production through OXPHOS. This was manifested as a decrease in both ATP generation and overall level of this nucleotide, vital for the efficient functioning of muscles having high energy demand [^28^]. Also, the observed decrease in the proportion of mitochondrial and glycolysis-derived ATP indicates that MVI deficiency significantly affects overall carbon metabolism, encompassing OXPHOS and glycolysis. Regarding the muscle, the most substantial change in intracellular ATP level was observed in MVI-KO SOL muscle, aligning with the reports showing that slow-twitch type fibers have ∼7-fold higher mitochondrial volume density and higher respiratory capacity compared to fast-twitch fibers [^27, 30^]. Since SOL muscle has high MVI expression, this observation further supports the link between MVI and mitochondria function.

AMPK maintains energy stores by regulating anabolic and catabolic pathways and therefore is considered as a central sensor of intracellular mitochondria status [^24^]. The AMPK activity is enhanced by energetic stress such as hypoxia, glucose deprivation, mitochondrial impairment and ATP depletion, evoking a series of biological events aimed at restoring the cellular metabolic balance between energy supply and demand. These events include the promotion of fatty acid oxidation and suppression of mTOR signaling [^23^]. AMPK activation is also involved in the fiber-type glycolytic-to-oxidative transition in skeletal muscle [^31,32^]. In MVI-KO muscles, we observed the AMPK activation and a decrease in glycolytic fibers (IIb, IIx) with a concomitant increase in oxidative fibers (I, IIa) in all examined muscles. These changes may be involved in the effective coupling of contraction and metabolism. Upregulation of AMPK activity in MVI-KO SOL muscle in all studied timepoints and downregulation of the mTOR pathway (i.e. a decrease of active forms of pp-70S6K, p-S6 and p-4E-BP1) indicate impairment in protein synthesis. Interestingly, these observations are tightly connected with our earlier findings showing a significant reduction in the cross-sectional area (CSA) of MVI-KO SOL and GM muscles with concomitant increase both in the average fiber number per selected area and in the number of smaller fibers [^2^]. These findings suggest the existence of a potentially novel muscle compensatory mechanism, the molecular mechanisms of which remain unknown. This adaptive response could aim at restoring perturbed energy homeostasis caused by the absence of MVI.

The above-described examination of the MVI-KO muscles at different developmental stages, including newborns, revealed that the observed differences between the muscles obtained from MVI-KO and WT mice result from the lack of MVI and not from a hyperactive phenotype characteristic for adult MVI-KO mice [^2,33^]. These findings underscore the important role of MVI in muscle metabolic homeostasis also during development.

Alterations in mitochondria functions in MVI-KO muscles imply that it could be also an important player in muscle lipid metabolism. This suggestion was verified *via* examination of the level and/or activity of the key enzymes involved in lipolysis, namely ATGL, HSL (conversely regulated by PKA and AMPK) and perilipin (PLN1) in MVI-KO SOL. PKA, phosphorylates HSL at Ser563, activating lipolysis [^34^]. It is known that AMPK regulates lipolysis in a complex way, often inhibiting hormone-sensitive lipase (HSL) by its phosphorylation at Ser565 thus preventing HSL activation by PKA. However, AMPK can also activate ATGL, stimulating basal lipolysis by breaking down triglycerides into fatty acids under energy-deficient conditions [^35^]. Such effect we observed in SOL muscle under the absence of MVI. This novel observation has been further confirmed by an elevated level of PLIN1, indicating that active HSL is more eager to associate with the droplet surface. Moreover, the observation that the level of ATGL, a major lipase governing lipolysis, is enhanced additionally confirms the involvement of MVI in lipolysis. The above-described changes in the level/activity of these lipases led to a decrease in the level of triacylglycerols (TAG) and a decrease in the mass of epidydimal fat pads. These observations are in line with the reports showing that ablation of ATGL significantly decreased lipolysis and fatty acids (FAs) release from lipid droplets [^36^]. Furthermore, a genome-wide meta-analysis highlighted the potential significance of the *MYO6* gene in the communication between the brain, adipose tissue, and liver in regulating lipid metabolism in humans [^37^].

Regarding the mechanisms behind the above-described observations, it is plasusible that one could be associated with the impairment of trafficking of proteins involved in muscle metabolism in the absence of MVI, especially since this motor is involved in endocytosis. Also, the disablement of an interaction of MVI with p-S6 (MVI-binding partner) under MVI-loss [^38,39^] could be responsible for the observed changes in protein synthesis. Furthermore, lack of MVI could affect organization of the mitochondrial network *via* the interaction with Parkin as it was shown in non-muscle cells [^9^]. Such effect was also reported for myo19, interacting with the outer mitochondrial membrane *via* its C-terminal tail domain (directly or indirectly by binding to mitochondrial proteins such as Miro1 and metaxins) and with actin filaments *via* the N-terminal motor domain [^40,41^]. Myo19 was also shown to tether mitochondria to the endoplasmic reticulum (ER) thus promoting mitochondria fission [^27^]. This observation is in line with our earlier results demonstrating that lack/depletion of MVI in myogenic cells affects the morphology of the ER, Golgi apparatus and adhesion complexes [^8,13^]. Furthermore, nuclear myosin 1 (NM1) has been implicated in mitochondrial biogenesis and function as it was shown that NM1 loss leads to reduced mitochondrial networks, underdeveloped inner membrane cristae, transcriptomic alterations in mitochondrial biogenesis, and a metabolic shift from OXPHOS to aerobic glycolysis [^42^]. Together, a growing line of evidence underscores the crucial roles of myosins in maintaining mitochondrial function and homeostasis.

Summarizing, we showed for the very first time that MVI plays an important role in muscle and myogenic cell energy metabolism. These novel observations indicate that MVI is involved in mitochondria respiration and ATP production as well as in modulation of protein synthesis, taking a part in “deciding” on the muscle fiber type composition during muscle growth and development, as well as in lipid metabolism. In our opinion, this work opens a new field both in the regulation of muscle energy and in myosin research.

## Acknowledgments

We thank Prof. Folma Buss from the Cambridge Institute for Medical Research, University of Cambridge, United Kingdom, for her gift of *SV* mice. Also, we thank Dr. Aleksandra Wyciszkewicz from Sartorius BioAnalytics for help in optimization of protocol regarding growth curves on glucose and galactose medium. Confocal and fluorescence imaging was performed at the Laboratory of Imaging Tissue Structure and Function which serves as an imaging core facility at the Nencki Institute of Experimental Biology and is a part of the infrastructure of the Polish Euro-BioImaging Node. This work was supported by the project financed by the Minister of Education and Science based on contract No 2022/WK/05 (Polish Euro-BioImaging Node “Advanced Light Microscopy Node Poland”).

## CRediT authorship contribution statement

**Dominika Wojton:** Conceptualization, Methodology, Investigation, Resources, Data Curation, Writing-Review&Editing, Visualization. **Dorota Dymkowska:** Methodology, Validation, Data Curation. **Damian Matysniak:** Methodology, Validation, Investigation. **Małgorzata Topolewska:** Investigation, Resources. **Maria Jolanta Redowicz:** Conceptualization, Validation, Data Curation, Writing - Review&Editing, Supervision, Project administration. **Lilya Lehka:** Conceptualization, Methodology, Validation, Formal analysis, Investigation, Resources, Data Curation, Writing – Original Draft, Writing – Review&Editing, Visualization, Supervision, Project administration, Funding acquisition.

## Data availability

The authors declare that any supporting data or material associated with this original research is available from corresponding authors under reasonable request.

