## Supplementary material for "Unconventional myosin VI is involved in regulation of muscle energy metabolism"

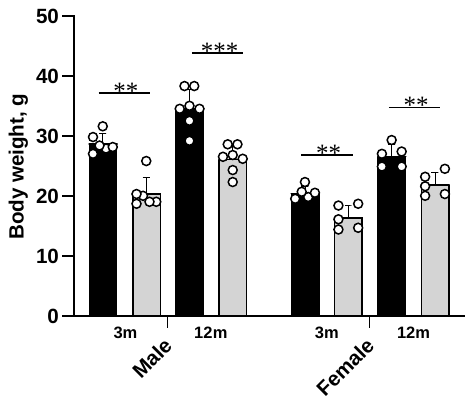

**Supplementary Fig 1.** **Analysis of the body weight in 3- and 12-month-old male and female WT and MVI-knockout mice.** Data are presented as mean value±SD. Statistical significance was analyzed with *t*-test, ***p* < 0.01 or ****p* < 0.001 related to WT animals in each age.

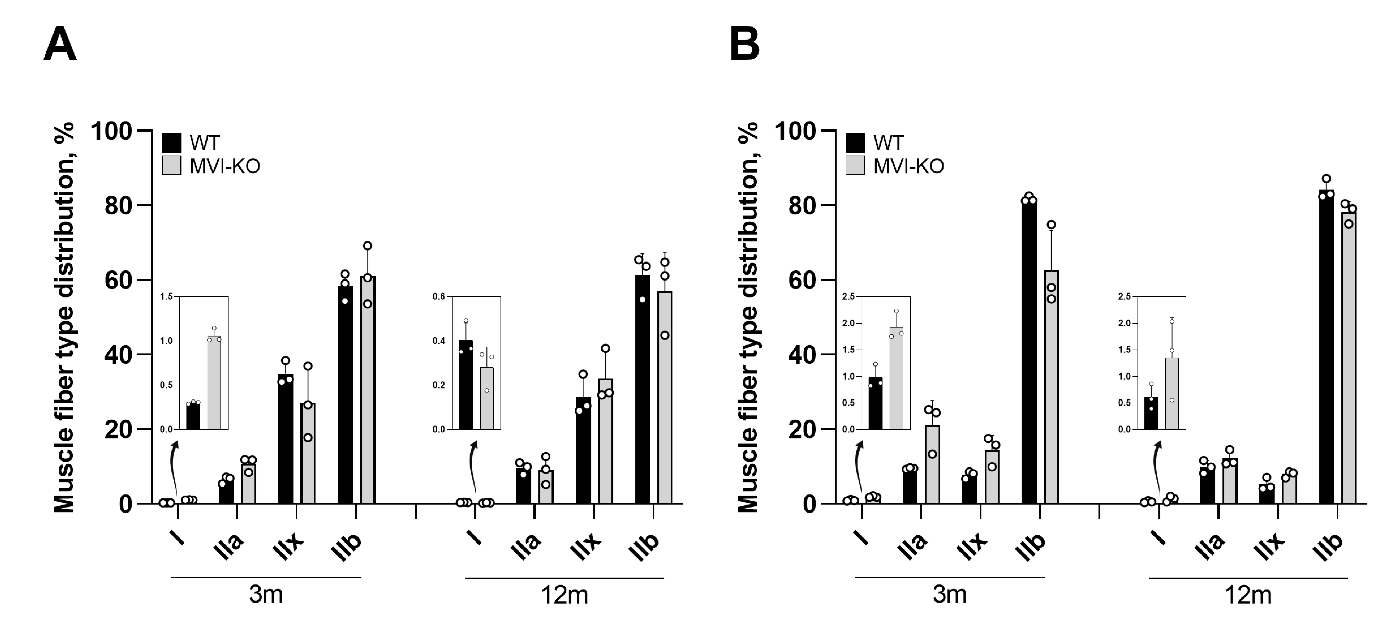

**Supplementary Fig 2.** **Glycolytic-to-oxidative fiber-type switch in myosin VI knockout (MVI-KO) muscles isolated from 3- (3m) and 12-months-old (12m) mice.** (A) Fiber typing *via* myosin heavy chain (MyHC) immunohistochemistry in MVI-KO and WT tibialis anterior (TA) muscle. Inserts, graphs representing the percentage of type I muscle fibers, pointed by arrows. (B) Fiber typing *via* MyHC immunohistochemistry in MVI-KO and WT gastrocnemius (GM) 3m and 12m muscle. Inserts, graphs representing the percentage of type I muscle fibers, pointed by arrow.

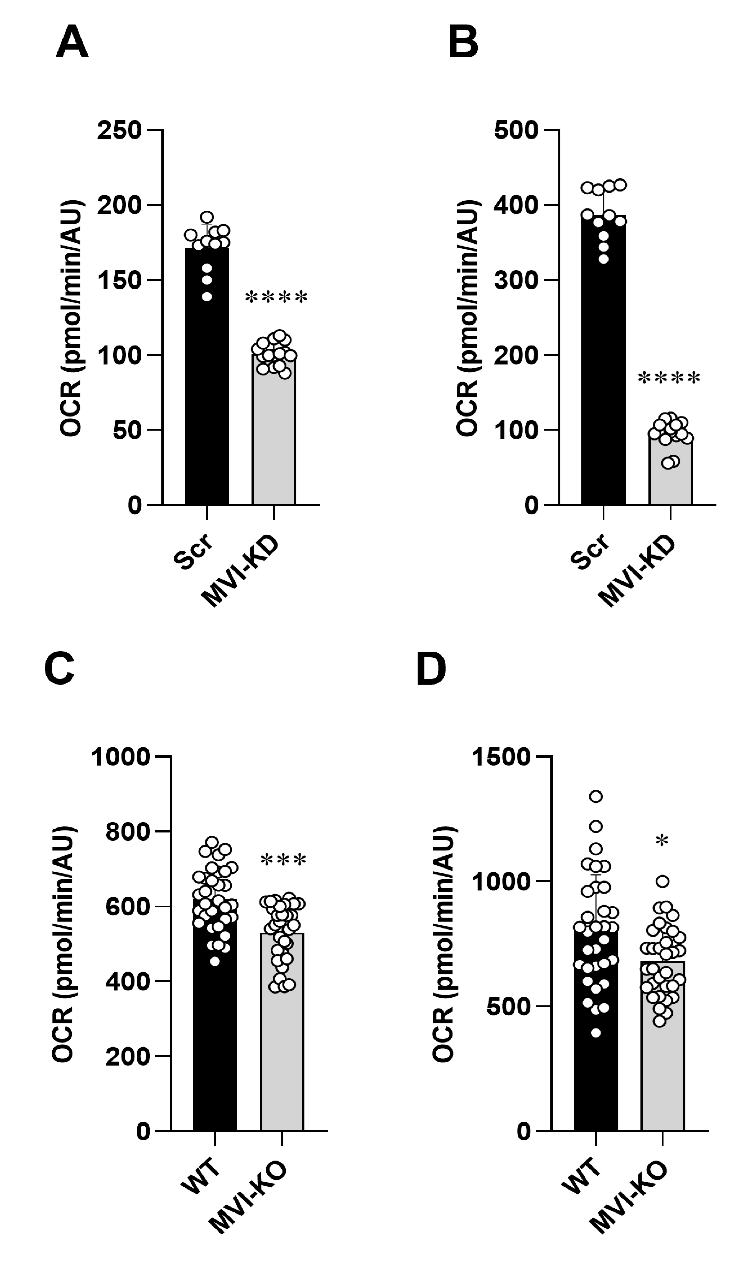

**Supplementary Fig 3. Mitochondrial respiratory control ratios in myogenic cells under myosin VI (MVI) deficit.** (A) basal respiration in control (Src) and MVI knockdown (MVI-KD) C2C12 cells; (B) maximal respiration in control (Src) and MVI knockdown (MVI-KD) C2C12 cells; (C) basal respiration in primary myoblasts isolated from muscles of heterozygous (WT) and MVI-knockout (MVI-KO) mice; (D) maximal respiration in primary myoblasts isolated from muscles of heterozygous (WT) and MVI-knockout (MVI-KO) mice. Data are represented as mean±SD, N = 3, n (technical replicates) ≥8. Statistical significance was analyzed with t-test, *p ≤0.05, ***p ≤0.001 and ****p ≤0.0001 vs. Scr) or WT cells.

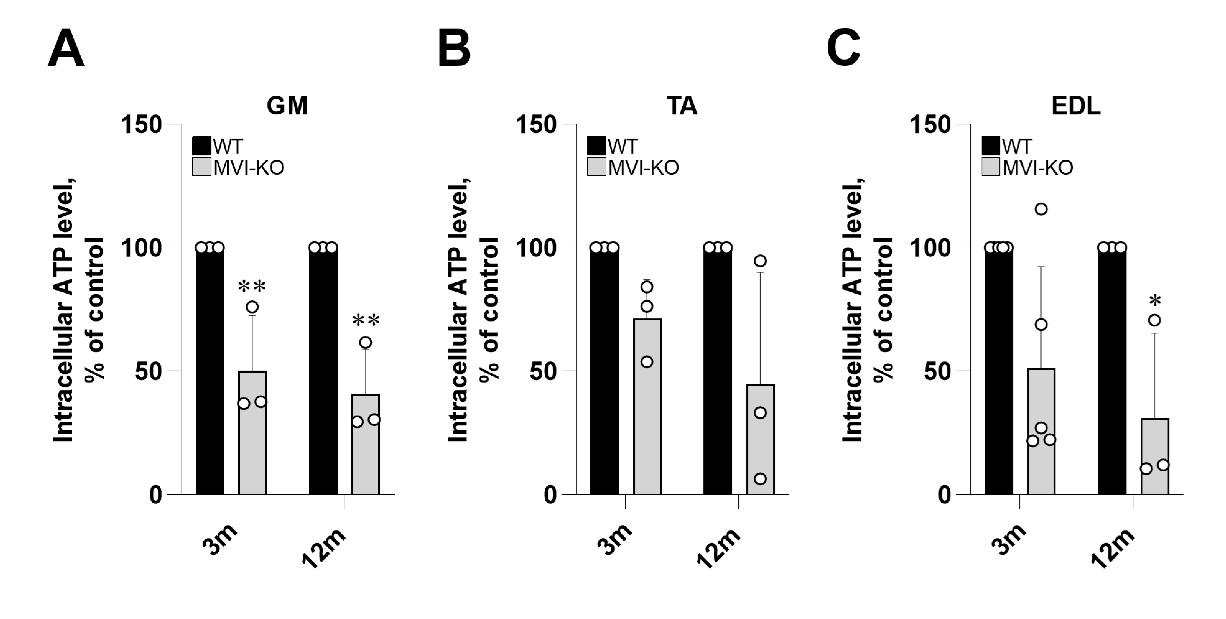

**Supplementary Fig 4. Reduction of ATP content in myosin VI knockout (MVI-KO) muscles.** (A) gastrocnemius (GM), (B) tibialis anterior (TA), and (C) extensor digitorum longus (EDL) muscles isolated from 3- (3m) and 12-months-old (12m) mice. Statistical significance was analyzed with t-test, *p <0.05, **p <0.01 vs. heterozygous (WT) muscles.

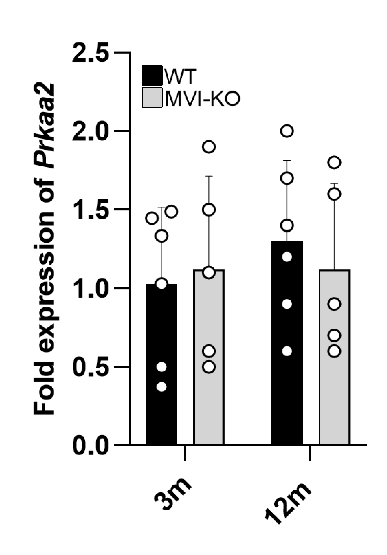
**Supplementary Fig 5. Relative expression of mRNA-*Prkaa2* (AMPK catalytic unit) in SOL of 3- and 12m animals.** mRNA-*B2m* (β-2 microglobulin) was used for normalization, 3m WT served as 1.
